# Thermal acclimation of stem respiration reduces global carbon burden

**DOI:** 10.1101/2024.02.23.581610

**Authors:** Han Zhang, Han Wang, Ian J. Wright, I. Colin Prentice, Sandy P. Harrison, Nicholas G. Smith, Andrea Westerband, Lucy Rowland, Lenka Plavcova, Hugh Morris, Peter B. Reich, Steven Jansen, Trevor Keenan

## Abstract

Stem respiration is a key driver of carbon flux from ecosystems to the atmosphere, yet its response to global warming remains poorly constrained. In particular it has been proposed that stem respiration acclimates to changing temperatures, which could have large implications for carbon cycling under climate change, but no theory exists to predict acclimated respiration rates. Here, we hypothesized that stem respiration is physiologically linked to transpiration in order to maintain hydraulic continuity. We then use that linkage, combined with Eco-evolutionary optimality theory, to develop a theoretical prediction of the temperature sensitivity of both acclimated and instantaneous stem respiration. Leveraging an extensive global dataset, we observe temperature sensitivities of stem respiration across geographical and seasonal variations that are consistent with this prediction. Our findings reveal that stem respiration contributes approximately a quarter of the global above-ground auto-trophic respiration, with an estimated annual emission of around 11.20 ± 5.88 Pg C—comparable to total anthropogenic emissions. Importantly, incorporating thermal acclimation of stem respiration into projections significantly reduces predicted land ecosystem carbon emissions by 4.41 and 9.56 Pg C under the SSP126 and SSP585 scenarios, respectively, for the 21st century.

---

Globally, plant respiration represents a significant proportion of annual carbon fluxes from terrestrial ecosystems to the atmosphere, and is approximately six times greater than anthropogenic emissions (Ciais et al., 2014; Friedlingstein et al., 2023). Plants respire half of their photosynthetic products to support eco-physiological processes (Campioli et al., 2016; Huntingford et al., 2017; Wang et al., 2020). As an enzyme-mediated process, at short time frames, respiration increases nearly exponentially with increasing temperature (Heskel et al., 2016). Additional respiratory carbon release is thus expected in a warming world, leading to more global warming within the next several decades (Cox et al., 2000; Huntingford et al., 2013).

However, plants can dynamically adjust the response of respiration to temperature over the long term (weeks to years), even though plant respiration always shows an accelerating increase when subjected to a short-term (minutes to hours) increase in temperature. Typically, a plant that has experienced warmer temperatures will have a lower rate of respiration at a standardized measurement temperature than a plant that has experienced cooler temperatures. This process is labeled thermal acclimation. The greater the thermal acclimation of respiration globally, the smaller the positive feedback between climate warming and ecosystem CO_2_ release (Reich et al., 2016). to long-term warming by reducing their reference respiration rates and/or reducing their thermal sensitivities (Gansert et al., 2002; Smith & Dukes 2013; Reich et al. 2016; Smith et al., 2019; Wang et al., 2020; Crous et al., 2022). This behavior could potentially weaken the positive feedback between climate warming and carbon release from plants (Atkin et al., 2003; Slot et al., 2014; Luo, 2007). Yet the mechanisms and the magnitude of this acclimation are still unclear. This is particularly true within the stems, which accounts for two thirds of above-ground plant biomass (Poorter et al. 2012; Zhou et al., 2021). Stem respiration is hypothesized to be more sensitive to environmental conditions than leaves or fine roots because leaves have multiple mechanisms for thermal regulation, and because soil temperatures fluctuate less than air temperature. Unfortunately, most studies of stem respiration focus on the local scale and there is a lack of studies globally and a theory-based understanding of stem respiration thermal acclimation (Lavigne et al., 1997; Rowland et al., 2018; Westerband et al., 2022).

The representation of respiration in Earth System Models (ESMs) is rather simplistic and largely untested. This introduces large uncertainties in predictions of the carbon-climate feedback (Lombardozzi et al., 2015; Collalti et al., 2020). For example, many ESMs do not explicitly implement the time-dependent acclimation of plant respiration to warming (Krinner et al., 2005; Niu et al., 2011). Those models that do consider acclimation assume that stem respiration has the same acclimation behavior as leaves or ignore it altogether (Clark et al., 2011; Lawrence et al., 2019; Thum et al., 2019, Butler et al., 2022), with highly empirical nature of all any approaches to incorporate stem respiration acclimation. An improved understanding of the magnitude of stem respiration and the mechanisms responsible for its acclimation is needed for reliable and robust predictions of the global carbon-climate feedback.

Eco-evolutionary optimality (EEO) principles have been shown to provide novel insights into, and parameter-sparse predictions of, plant eco-physiological processes (Harrison et al., 2021). Indeed, a recent EEO-based study (Ren et al., 2024) has shown that leaf respiration can be thought of as reflecting the costs associated with maintaining a given carboxylation capacity, such that its acclimation is largely driven by carboxylation capacity demand. This suggests that EEO principles might together provide a valid and more informative approach for investigating the controls on, and magnitude of, thermal acclimation of stem respiration.

The goal of this study is to develop theory of thermal acclimation of stem respiration from first principles and to test these theoretical predictions globally with multiple independent datasets. Specifically, we aim to answer the following questions: (1) What drives the thermal acclimation of stem respiration? (2) How does this thermal acclimation affect the global carbon cycle?

Our demand-driven theory is based on the hypothesis that stem respiration occurs to maintain the hydraulic continuum, as determined by the transpiration rates, and therefore demand for water within the canopy. The energy released from stem respiration supports hydraulic continuum, as a proxy of associated processes, such as storage capacity, producing new xylem and the maintenance of stem water potential (Martorell et al., 2014; Zeppel et al., 2019; Tomasella et al., 2020; Gauthey et al., 2022). Guiding by Eco-evolutionary optimality theory, this coupling ensures that stem respiration is neither more--which would waste additional, non-productive carbon consumption--nor less than what is required for supporting the hydraulic continuum--which would threat the hydraulic security. We combined this hypothesis with physical principles to quantitatively predict the spatial and temporal thermal acclimation of stem respiration. We first assumed that whole-plant stem respiration (*R*_s_, μmol C m^−2^ s^−1^) is proportional to canopy transpiration demand (E, μmol H_2_O m^−2^ s^−1^) with a cost factor representing the respiration rate required to support a unit transpiration rate. In principle, this cost factor should be influenced by temperature: water is less viscous as temperature increases and thus the cost incurred for supporting a given transpiration rate is lower. Given that *R*_s_ is a product of the mass-based respiration rate over time (*r*_s_, μmol C g^−1^ s^−1^) and total biomass (*M*_s_, g C), we expect that *r*_s_ and *M*_s_ also coordinate with *E* and the cost factor, proportionally. Then we captured the leading variable in different scales to abstract the theory. At a global scale, where spatial variation in both *r*_s_ and *M*_s_ is potentially substantial, we assumed that *r*_s_ and *M*_s_ track variation in the cost factor and *E*, respectively. Therefore, the thermal response of the cost factor as determined by water viscosity mainly controls the variation in *r*_s_ along the temperature gradient, globally. For temporal changes at a weekly to seasonal scale, where the change in *M*_s_ is relatively marginal compared to the changes in transpiration demand, we expect only *r*_s_ to be coordinated with temporal variation in both *E* and the cost factor. Therefore, the response of *r*_s_ to temperature that emerges from temporal changes should be the same as that which emerges from the spatial patterning when its response to *E* is also taken into account.

To generate empirically testable quantitative predictions for how stem respiration adjusts to temperature, we chose two forms of respiration rate to characterize this ability. One is r_s_ at growth temperature (r_s.gt_, μmol C g^−1^ s^−1^), which is the respiration rate under typical temperature conditions in habitats, the other one is r_s_ at a common temperature of 25°C (r_s25_, μmol C g^−1^ s^−1^), which is the promise for unified comparison of respiratory capacity (and associated investment in mitochondrial protein) to be made among contrasting sites and species (Atkin et al., 2015). Specifically, we predicted that rs at growth temperature acclimates by declining of temperature by 2.3 % K^−1^. When evaluating r_s_ at 25° C, an even steeper decline, of 10 % K^−1^, would be expected due to the immediate response of respiration to temperature, that is Arrhenius relationship between r_s_ and temperature (Figure 1). This response is opposite in sign to the instantaneous response only derived from enzyme kinetics (+7.88% K^−1^).

**Fig 1.**
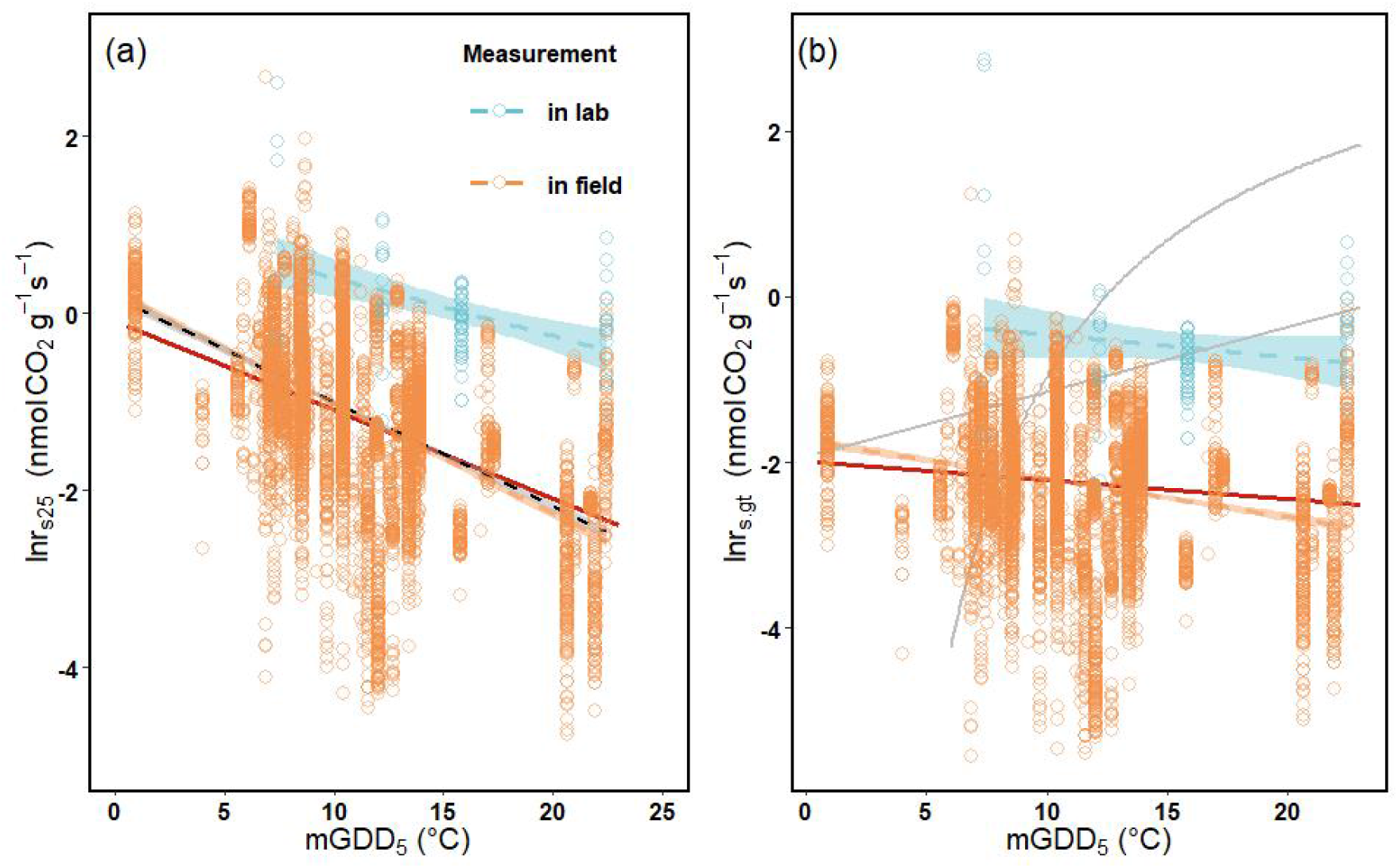
Global trends of stem respiration at reference (r_s25_) and at growing temperature (r_s.gt_) in relation to the mean growing season temperature (mGDD_5_). **(a)** stem respiration was measured in the laboratory (blue circles) and in the field (orange circles). r_s25_ represents stem respiration standardized to the reference temperature of 25 °C, and mGDD_5_ is the mean temperature of the growing season defined as days when the temperature is >5°C. Dashed fitted lines are shown for the laboratory data (blue line), field data (orange line) and all data (black line) separately, after excluding two outliers with values >3 (Mathematical details are provided in Table 1). The shaded area represents the 95% confidence intervals. The solid red line is the theoretical prediction of thermal sensitivity, with a coefficient of −0.1. **(b)** r_s.gt_ represents stem respiration standardized to the mean growing temperature, using the same colors as in (a). The solid red line is the theoretical prediction of −0.023. The instantaneous thermal response was characterized by the Arrhenius equation (gray non-linear curve) and a fixed-Q_10_ (gray line).

This thermal response is well confirmed by empirical analysis using Global Stem Respiration Dataset (GSRD), which includes 212 species sampled at 96 field sites worldwide spanning all vegetated continents, climate zones, and major plant functional types (Figure S1). A linear regression of *r*_s_ against mean growing season temperature mGDD_5_ showed a strong negative response of *r*_s_ to warming across all the GSRD sites: *r*_s25_ decreased by 11.8 ± 0.3 % K^−1^ warming, consistent with, but slightly stronger than, our theoretical prediction (–10% K^−1^) (Figure 1a, Table 1). The *in field* and *in lab* measurements showed a similar thermal response in *r*_s25_ (field measurements: –12.5 ± 0.3 % K^−1^; laboratory measurements: –10.6 ± 0.2 % K^−1^). The absolute value of the *in lab* measurements were generally larger, possibly because some CO_2_ leaks into upward sap flow in the field, and because CO_2_ diffusion limitations from bark are removed. Comparisons using site-mean values, to minimize the potential impact of over-representation of individual sites with much larger sample sizes, showed that *r*_s25_ decreased by 8.3 ± 2.1 % K^−1^ warming (Figure S2, Table S1). We also tested whether using the Arrhenius equation to calculate *r*_s25_ impacted the response using only sites where respiration rate was directly measured at a temperature of 25 ± 1°C; these data showed a similar thermal response in *r*_s25_ (site-mean: 8.0 ± 3.0% K^−1^, all individuals: 13.4 ± 0.8% K^−1^) (Figure S3, Table S2). Thus, considered multiple ways, the observed response of *r*_s25_ against mGDD_5_ falls within the range of ca. –8% K^−1^ to –13% K^−1^. Considering next stem respiration under typical growing conditions (Figure 1b and Table 1), both observations and theory show a significant negative response of *r*_s.gt_ to temperature (data-based: –3.9 ± 0.3% K^−1^, theory-based: –2.26% K^−1^),), contrary in sign to the traditional scheme only considering enzyme kinetics. In contrast to leaf respiration, the stem respiration declares a much stronger thermal acclimation, while leaf respiration generally shows a weak but still positive response to warming (Wang et al., 2020). The difference in the response of stem and leaf respiration inspired us, when Earth System Models (ESM) routinely begin to account for acclimation, they need to do it differently for stem wood, leaves and roots, however by now, most still ignore it entirely.

**Table 1.** Statistical details of the regression analysis. r_s25_ represents stem respiration standardized to the reference temperature of 25 °C. r_s.gt_ represents stem respiration standardized to the mean growing temperature mGDD_5_, which is the mean temperature of the growing season defined as days when the temperature is >5°C. These are statistical details of Figure 1, showing the relationship between r_s25_/r_s.gt_ and mGDD_5_.

| Quantity | Theoretical prediction | Measurements | fitted coefficient | confidence intervals | | Intercept<br>(mean ± SE) | $R^2$ | p | df |
| --- | --- | --- | --- | --- | --- | --- | --- | --- | --- |
|  |  |  |  | 2.50% | 97.50% |  |  |  |  |
| $r_{s25}$ | -0.1 | In lab | -0.106 | -0.126 | -0.086 | $1.79 \pm 0.33$ | 0.25 | <0.001 | 79 |
| | | In field | -0.125 | -0.128 | -0.122 | $0.23 \pm 0.03$ | 0.28 | <0.001 | 5194 |
| | | All data | -0.118 | -0.121 | -0.115 | $0.19 \pm 0.04$ | 0.25 | <0.001 | 5275 |
| $r_{s,gt}$ | -0.023 | In lab | -0.027 | -0.047 | -0.007 | $-0.18 \pm 0.33$ | 0.01 | 0.18 | 79 |
| | | In field | -0.046 | -0.049 | -0.043 | $-1.74 \pm 0.03$ | 0.05 | <0.001 | 5194 |
| | | All data | -0.039 | -0.042 | -0.037 | $-1.79 \pm 0.04$ | 0.04 | <0.001 | 5275 |

The boreal site (55.9°N, 98.75°W) in the GSRD provides continuous data on *r*_s_ through the growing season. We estimated the daily transpiration demand using daily radiation, temperature, vapor pressure deficit and air pressure inputs to an EEO-based evapotranspiration model (Tan et al., 2021, 2023). Multiple regression of log-transformed *r*_s25_ against the mean values of air temperature and log-transformed transpiration showed a significant negative response of *r*_s25_ to temperature (–6.8 ± 0.5% K^−1^ and –10.4 ± 0.6% K^−1^ for the two temperature time-scales, separately) and a positive response to transpiration (92 ± 2.9 % and 106 ± 3.2 % for the two temperature time-scales, separately). This is consistent with the theoretical predictions (–10 % K^−1^ and 100 % for temperature and transpiration, respectively) (Figure 2); the small differences between these values likely reflect uncertainties in the two predictors or the influence of other potential explanatory variables such as soil moisture (Huang et al., 2017; Cui et al., 2018). The warming experiment provides further information on the magnitude of thermal acclimation at a weekly timescale. The five species showed a consistent decrease in *r*_s25_ with warming with an overall trend of 9.0 ± 0.7% K^−1^, close to our theoretical prediction of 10% (Figure 3), and similar downward trends were found for each species (Table S3). A comparison of the four *Pinus* species showed a somewhat more muted thermal acclimation response than that obtained for *Pinus* in the GSRD compilation (–15.0 ± 1.0 % K^−1^), possibly reflecting differences in the temperature range sampled (Figure S4 and Table S4).

**Fig 2.**
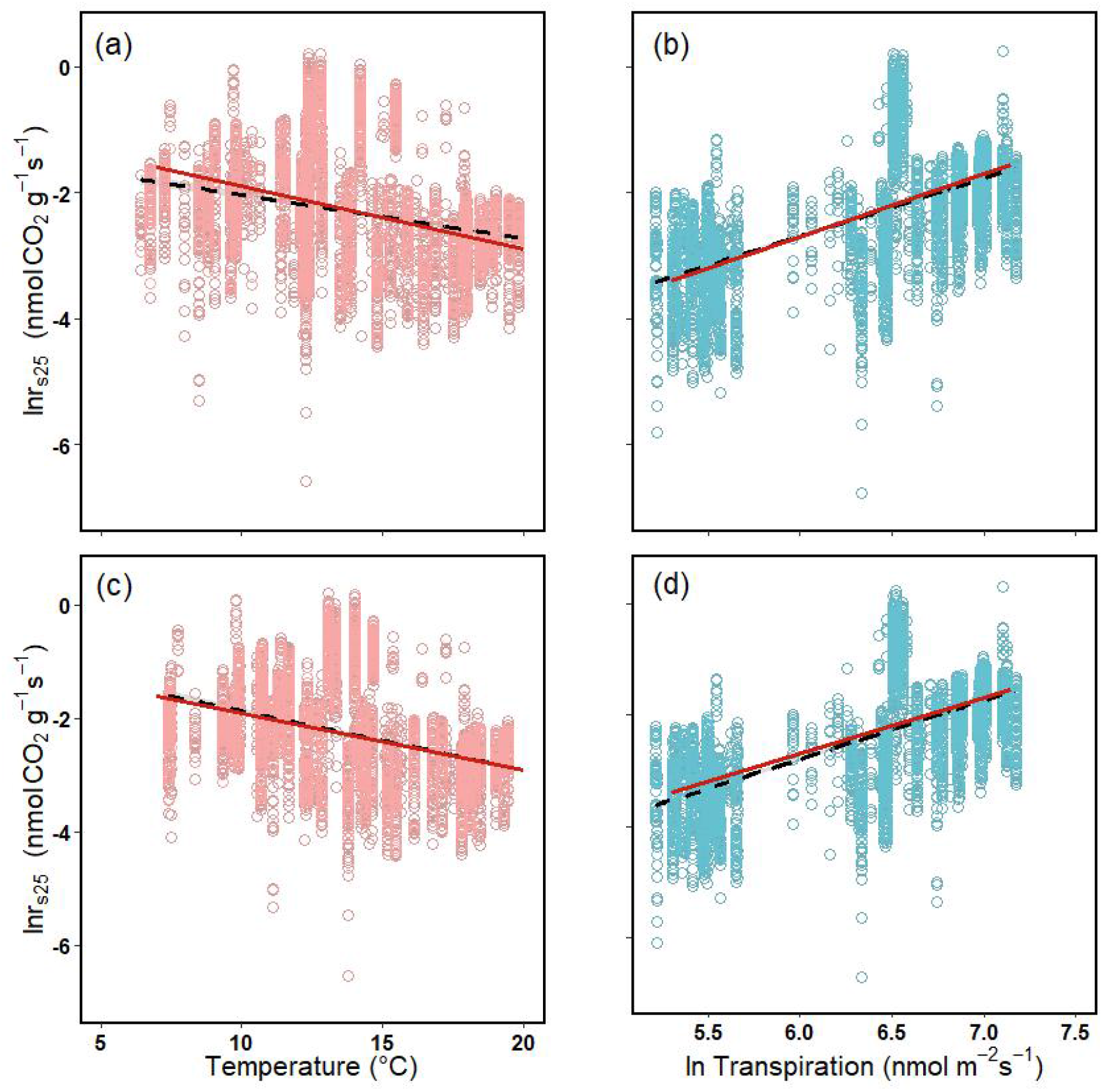
Effect of temperature and transpiration on stem respiration at seasonal scale. **(a)** Temperature is the mean value of the three days before stem respiration was measured. r_s25_ is stem respiration, natural-log transformed, at the reference temperature of 25°C. **(b)** Transpiration is calculated over the seven days before measurement (see Methods) and was natural-log transformed. The dashed line is the relationship fitted by multiple regression (lnr_s25_ = −0.068 ± 0.005*Temperature+0.92 ± 0.029* lnTranspiration, R^2^=0.22, VIF=1.83). The shaded area represents the 95% confidence intervals. **(c) and (d)** are the same as (a) and (b), except that the temperature was averaged over four days instead of three. The dashed line shows the fitted relationship by multiple regression (lnr_s25_ = −0.104 ± 0.006*Temperature+1.06 ± 0.032 * lnTranspiration, R^2^=0.23, VIF=2.32).

**Fig 3.**
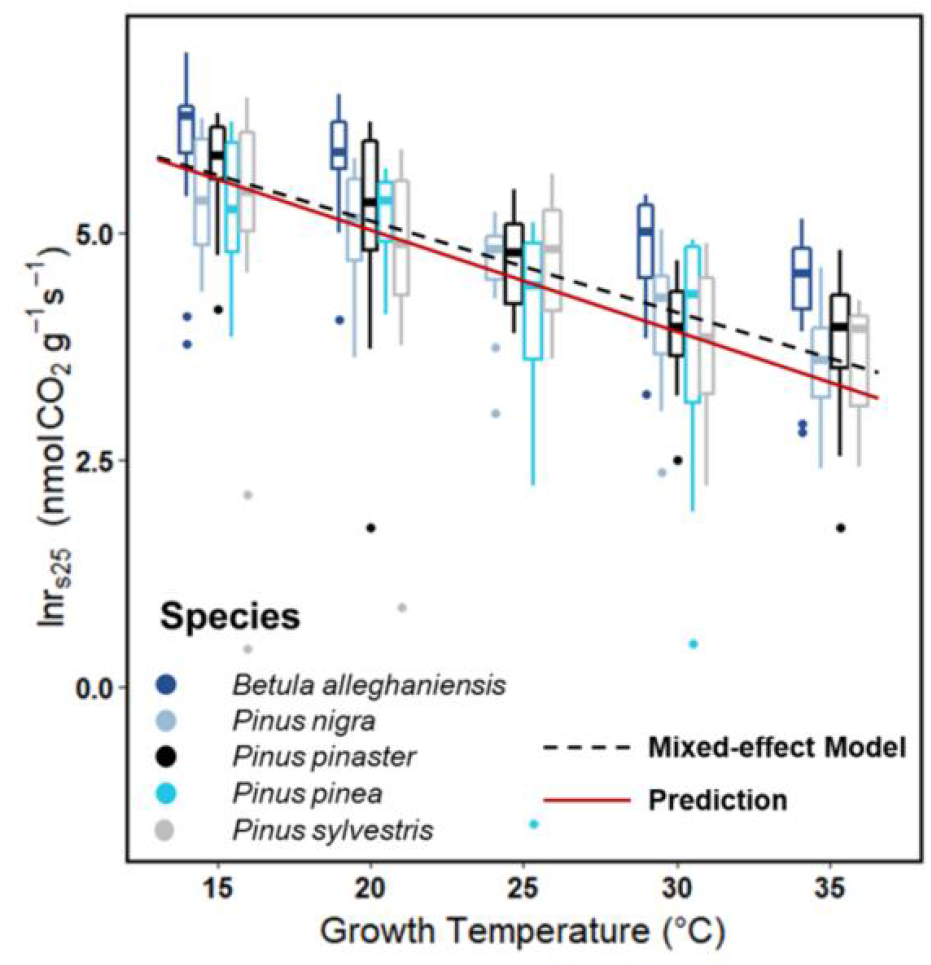
Thermal sensitivity of stem respiration derived from a warming experiment. “Growth Temperature” is the temperature at which individual species were grown. The warming experiment includes five species, and the fitted lines used here are based on the data from all samples. Boxplots represent different replicate groups of the same species at the same environmental temperature; the black vertical lines show the mean values and the boxes show the interquartile range The dashed line is fitted (lnr_s25_ = −0.0896±0.007 Temperature, R^2^=0.37) using a linear mixed-effects model, where species was a random effect. The red solid line is our theoretical prediction.

The limited nature of the observations of stem respiration, whether based on in field measurements or laboratory experiments, introduces some uncertainties in our analysis. In particular, we made assumptions about mean diameter values and living cells mass to derive mass-based estimates of respiration. Furthermore, there is a bias in the sampling, with relatively poor representation of the tropics and southern hemisphere. Nevertheless, the congruence of the results obtained from different sources and subsets of the data (Figure 1a, Figure 1b, Figure S2, Figure S3) and, more importantly, the similarity between the observed and predicted relationships suggests that our conclusions about the thermal response of stem respiration is robust. However, these data limitations currently preclude any analysis of the sensitivity of respiration to other factors, such as VPD or CO_2_

Based on the agreement between theoretical and empirical predictions, we developed a simple model to estimate the global carbon flux from stem respiration and the impact of acclimation on the global carbon cycle in the future. In this model, the total stem respiration rate per unit area is predicted as the product of stem biomass and *r*_s.gt_ on an annual time-step. The amount of respiring biomass is estimated from total above-ground stem biomass and parenchyma tissue fraction, which was approximated and simulated as a function of growth temperature (Morris et al., 2016). The information on above-ground stem biomass was at a resolution of 1 ha (Santoro et al., 2021). Mean annual temperature during the growing season, defined as the interval when temperatures were > 5°C, was used to represent the annual growth thermal environment for woody species. *r*_s.gt_ was estimated for each grid by temperature conversion with a parameter by global parameter calibration. According to this model, the annual CO_2_ released globally by stem respiration was 11.20 ± 5.88 Pg C around 2010 (Figure 4a). This is comparable to annual anthropogenic carbon emissions (9.6 ± 0.5 Pg C: Friedlingstein et al., 2023), but is only about one third of the annual respiration flux released from leaves, which has been estimated as about 30 Pg C (Atkin et al., 2007). Thus, global variation in annual stem respiration appears to be largely controlled by stem biomass: regions with higher biomass have higher respiration. Consequently, high stem respiration was not only found in tropical forests, but also in the humid temperate forests (for instance, in eastern North America, eastern Asia, and Europe).

**Fig 4.**
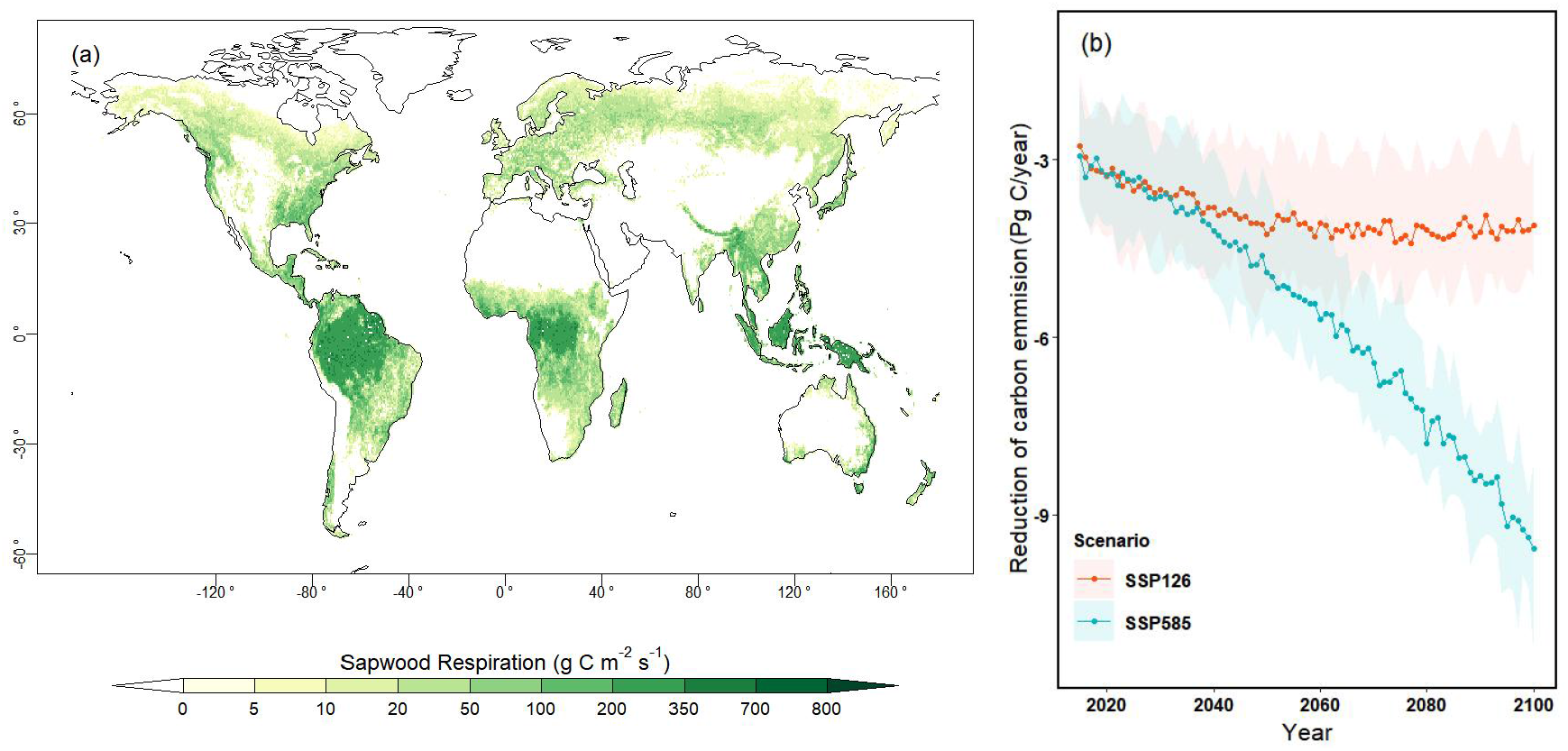
Predicted global stem respiration and a simulation of its future response. **(a)** stem respiration is calculated at the global scale using a gridded system, at modern levels of CO_2_ emissions (see details in supplementary 3.2 Modeling and Simulation). **(b)** A simulated reduction in carbon emissions was generated by considering thermal acclimation responses over the 21st century, where the temperature was derived from an ensemble of four future climate simulations (CABLE, ORCHIDEE, JULES and CLM5). The dashed dotted line is the mean ensemble value (orange: under SSP126, blue: under SSP585) and the shaded area represents the range of the model ensemble.

Finally, we applied this modelling scheme using climate change scenarios to assess the potential influence of thermal acclimation on the carbon flux from stem respiration over the 21^st^ century. Our results show that implementing thermal acclimation in models substantially reduces predicted land ecosystem carbon emissions by 4.41 and 9.56 Pg C by 2100 under the different human behaviors -- SSP126 (low emissions) and SSP585 (high emissions) scenarios, respectively (Figure 4b). The negative feedback of thermal acclimation to climate is more pronounced in scenarios with greater warming. We compared our results with outputs from the Community Land Model 5.0 (CLM5), which takes account of the thermal acclimation of plant respiration. In CLM5, *r*_s_ is predicted to decrease with temperature by 1.83 % K^−1^, which is an order of magnitude less than shown by our empirical analyses. This underestimate of the thermal acclimation of stem respiration in CLM5 leads to a higher carbon emission from terrestrial ecosystems, resulting in an overestimation of 3.6 and 7.8 Pg C in 2100 under the SSP126 and SSP585 scenarios, respectively (Figure S7).

We have demonstrated that the application of EEO principles provides a promising approach to predicting stem respiration and its thermal acclimation in a simple way that is consistent with empirical evidence. Specifically, our theoretical model links stem respiration as represented by the carbon cost of maintaining effective hydraulics together with leaf respiration for maintaining carboxylation, and constrains the competition between these two costs via optimal stomatal behaviour. This approach could be implemented in ESMs to improve the accuracy of these models. Moreover, it is important to do this because, although carbon released by stem respiration is one third as much as that from leaf respiration, it shows a much stronger acclimation to temperature. This suggests a potential for enhanced carbon use efficiency in a warming world, a subject on which there has been considerable debate (Vicca et al., 2012; Campioli et al., 2015; Manzoni et al., 2018). Enhanced carbon use efficiency would, in turn, substantially weaken the currently assumed positive climate-carbon feedback due to the acclimation of respiration, or even lead to negative climate-carbon feedback.

## Author contributions

Theory: H.Z., H.W., I.C.P., I.J.W., and S.P.H. Data: H.Z., I.J.W., A.W., N.S., P.R., L.R., H.M., L.P. and S.J. Investigation: H.Z., H.W., I.C.P., S.P.H., I.J.W. and T.K. Visualization: H.Z. Writing: original draft: H.Z., H.W., S.P.H. Writing review and editing: all.

## Data and materials availability

All data needed to evaluate the conclusions in the paper are in the Supplementary Materials.

## References

1. A. Collalti, A. Ibrom, A. Stockmarr, A. Cescatti, R. Alkama, M. Fernández-Martínez, G. Matteucci, S. Sitch, P. Friedlingstein, P. Ciais, D. S. Goll, J. E. M. S. Nabel, J. Pongratz, A. Arneth, V. Haverd, I. C. Prentice, Forest production efficiency increases with growth temperature. Nat. Commun. 11, 5322 (2020).

2. A. C. Westerband, I. J. Wright, A. S. D. Eller, L. A. Cernusak, P. B. Reich, O. Perez-Priego, S. S. Chhajed, L. B. Hutley, C. E. R. Lehmann, Nitrogen concentration and physical properties are key drivers of woody tissue respiration. Ann Bot-london 129, 633 – 646 (2022).

3. A. Gauthey, J. M. R. Peters, R. Lòpez, M. R. Carins-Murphy, C. M. Rodriguez-Dominguez, D. T. Tissue, B. E. Medlyn, T. J. Brodribb, B. Choat, Mechanisms of xylem hydraulic recovery after drought in Eucalyptus saligna. Plant Cell Environ 45, 1216–1228 (2022).

4. C. Huntingford, P. Zelazowski, D. Galbraith, L. M. Mercado, S. Sitch, R. Fisher, M. Lomas, A. P. Walker, C. D. Jones, B. B. B. Booth, Y. Malhi, D. Hemming, G. Kay, P. Good, S. L. Lewis, O. L. Phillips, O. K. Atkin, J. Lloyd, E. Gloor, J. Zaragoza-Castells, P. Meir, R. Betts, P. P. Harris, C. Nobre, J. Marengo, P. M. Cox, Simulated resilience of tropical rainforests to CO2-induced climate change. Nat. Geosci. 6, 268–273 (2013).

5. C. Huang, J. Domec, E. J. Ward, T. Duman, G. Manoli, A. J. Parolari, G. G. Katul, The effect of plant water storage on water fluxes within the coupled soil–plant system. N. Phytol. 213, 1093–1106 (2017).

6. C. Huntingford, O. K. Atkin, A. M. la Torre, L. M. Mercado, M. A. Heskel, A. B. Harper, K. J. Bloomfield, O. S. O’Sullivan, P. B. Reich, K. R. Wythers, E. E. Butler, M. Chen, K. L. Griffin, P. Meir, M. G. Tjoelker, M. H. Turnbull, S. Sitch, A. Wiltshire, Y. Malhi, Implications of improved representations of plant respiration in a changing climate. Nat. Commun. 8, 1602 (2017).

7. Z. Cui, D. Xu, Z. Yang, N. Zhang, Z. Hong, Effects of soil moisture content on stem respiration rate, stem growth and sapwood nitrogen content in Dalbergia odorifera. Journal of South China Agricultural University. 39(2): 54–61 (2018).

8. D. B. Clark, L. M. Mercado, S. Sitch, C. D. Jones, N. Gedney, M. J. Best, M. Pryor, G. G. Rooney, R. L. H. Essery, E. Blyth, O. Boucher, R. J. Harding, P. M. Cox, The Joint UK Land Environment Simulator (JULES), Model description - Part 2: Carbon fluxes and vegetation. doi: 10.5194/gmdd-4-641-2011 (2011).

9. D. Gansert, K. Backes, T. Ozaki, Y. Kakubari, Seasonal variation of branch respiration of a treeline forming (Betula ermanii Cham.) and a montane (Fagus crenata Blume) deciduous broad-leaved tree species on Mt. Fuji, Japan. Flora - Morphol., Distrib., Funct. Ecol. Plants 197, 186–202 (2002).

10. D. L. Lombardozzi, G. B. Bonan, N. G. Smith, J. S. Dukes, R. A. Fisher, Temperature acclimation of photosynthesis and respiration: A key uncertainty in the carbon cycle-climate feedback. Geophys Res Lett 42, 8624–8631 (2015).

11. D. M. Lawrence, R. A. Fisher, C. D. Koven, K. W. Oleson, S. C. Swenson, G. Bonan, N. Collier, B. Ghimire, L. Kampenhout, D. Kennedy, E. Kluzek, P. J. Lawrence, F. Li, H. Li, D. Lombardozzi, W. J. Riley, W. J. Sacks, M. Shi, M. Vertenstein, W. R. Wieder, C. Xu, A. A. Ali, A. M. Badger, G. Bisht, M. Broeke, M. A. Brunke, S. P. Burns, J. Buzan, M. Clark, A. Craig, K. Dahlin, B. Drewniak, J. B. Fisher, M. Flanner, A. M. Fox, P. Gentine, F. Hoffman, G. Keppel-Aleks, R. Knox, S. Kumar, J. Lenaerts, L. R. Leung, W. H. Lipscomb, Y. Lu, A. Pandey, J. D. Pelletier, J. Perket, J. T. Randerson, D. M. Ricciuto, B. M. Sanderson, A. Slater, Z. M. Subin, J. Tang, R. Q. Thomas, M. V. Martin, X. Zeng, The Community Land Model Version 5: Description of New Features, Benchmarking, and Impact of Forcing Uncertainty. J Adv Model Earth Sy 11, 4245–4287 (2019).

12. G. Krinner, N. Viovy, N. de Noblet-Ducoudré, J. Ogée, J. Polcher, P. Friedlingstein, P. Ciais, S. Sitch, I. C. Prentice, A dynamic global vegetation model for studies of the coupled atmosphere-biosphere system. Glob. Biogeochem. Cycles 19 (2005).

13. G. Niu, Z. Yang, K. E. Mitchell, F. Chen, M. B. Ek, M. Barlage, A. Kumar, K. Manning, D. Niyogi, E. Rosero, M. Tewari, Y. Xia, The community Noah land surface model with multiparameterization options (Noah-MP): 1. Model description and evaluation with local-scale measurements. J. Geophys. Res.: Atmos. 116 (2011).

14. H. Wang, O. K. Atkin, T. F. Keenan, N. G. Smith, I. J. Wright, K. J. Bloomfield, J. Kattge, P. B. Reich, I. C. Prentice, Acclimation of leaf respiration consistent with optimal photosynthetic capacity. Global Change Biol 26, 2573–2583 (2020).

15. H. Morris, L. Plavcová, P. Cvecko, E. Fichtler, M. A. F. Gillingham, H. I. Martínez-Cabrera, D. J. McGlinn, E. Wheeler, J. Zheng, K. Ziemińska, S. Jansen, A global analysis of parenchyma tissue fractions in secondary xylem of seed plants. New Phytol 209, 1553–1565 (2016).

16. H. Poorter, K. J. Niklas, P. B. Reich, J. Oleksyn, P. Poot, L. Mommer, Biomass allocation to leaves, stems and roots: meta-analyses of interspecific variation and environmental control. N. Phytol. 193, 30–50 (2012).

17. J. R. A. Butler, R. M. Wise, S. Meharg, N. Peterson, E. L. Bohensky, G. Lipsett-Moore, T. D. Skewes, D. Hayes, M. Fischer, P. Dunstan, ‘Walking along with development’: Climate resilient pathways for political resource curses. Environ. Sci. Polic. 128, 228–241 (2022).

18. K. Y. Crous, J. Uddling, M. G. D. Kauwe, Temperature responses of photosynthesis and respiration in evergreen trees from boreal to tropical latitudes. N. Phytol. 234, 353 – 374 (2022).

19. L. Rowland, A. C. L. da Costa, A. A. R. Oliveira, R. S. Oliveira, P. L. Bittencourt, P. B. Costa, A. L. Giles, A. I. Sosa, I. Coughlin, J. L. Godlee, S. S. Vasconcelos, J. A. S. Junior, L. V. Ferreira, M. Mencuccini, P. Meir, Drought stress and tree size determine stem CO2 efflux in a tropical forest. N. Phytol. 218, 1393–1405 (2018).

20. M. Santoro, O. Cartus, N. Carvalhais, D. M. A. Rozendaal, V. Avitabile, A. Araza, S. de Bruin, M. Herold, S. Quegan, P. Rodríguez-Veiga, H. Balzter, J. Carreiras, D. Schepaschenko, M. Korets, M. Shimada, T. Itoh, Á. M. Martínez, J. Cavlovic, R. C. Gatti, P. da C. Bispo, N. Dewnath, N. Labrière, J. Liang, J. Lindsell, E. T. A. Mitchard, A. Morel, A. M. P. Pascagaza, C. M. Ryan, F. Slik, G. V. Laurin, H. Verbeeck, A. Wijaya, S. Willcock, The global forest above-ground biomass pool for 2010 estimated from high-resolution satellite observations. Earth Syst. Sci. Data 13, 3927–3950 (2021).

21. M. A. Heskel, O. S. O’ Sullivan, P. B. Reich, M. G. Tjoelker, L. K. Weerasinghe, A. Penillard, J. J. G. Egerton, D. Creek, K. J. Bloomfield, J. Xiang, F. Sinca, Z. R. Stangl, A. M. la Torre, K. L. Griffin, C. Huntingford, V. Hurry, P. Meir, M. H. Turnbull, O. K. Atkin, Convergence in the temperature response of leaf respiration across biomes and plant functional types. Proc National Acad Sci 113, 3832–3837 (2016).

22. M. B. Lavigne, M. G. Ryan, Growth and maintenance respiration rates of aspen, black spruce and jack pine stems at northern and southern BOREAS sites. Tree Physiol. 17, 543 –551 (1997).

23. M. Campioli, S. Vicca, S. Luyssaert, J. Bilcke, E. Ceschia, F. S. C. III, P. Ciais, M. Fernández-Martínez, Y. Malhi, M. Obersteiner, D. Olefeldt, D. Papale, S. L. Piao, J. Peñuelas, P. F. Sullivan, X. Wang, T. Zenone, I. A. Janssens, Biomass production efficiency controlled by management in temperate and boreal ecosystems. Nat. Geosci. 8, 843 –846 (2015).

24. M. Campioli, Y. Malhi, S. Vicca, S. Luyssaert, D. Papale, J. Peñuelas, M. Reichstein, M. Migliavacca, M. A. Arain, I. A. Janssens, Evaluating the convergence between eddy-covariance and biometric methods for assessing carbon budgets of forests. Nat. Commun. 7, 13717 (2016).

25. M. Slot, K. Kitajima, General patterns of acclimation of leaf respiration to elevated temperatures across biomes and plant types. Oecologia 177, 885–900 (2015).

26. M. Tomasella, E. Petrussa, F. Petruzzellis, A. Nardini, V. Casolo, The Possible Role of Non-Structural Carbohydrates in the Regulation of Tree Hydraulics. Int. J. Mol. Sci. 21, 144 (2019).

27. M. J. B. Zeppel, W. R. L. Anderegg, H. D. Adams, P. Hudson, A. Cook, R. Rumman, D. Eamus, D. T. Tissue, S. W. Pacala, Embolism recovery strategies and nocturnal water loss across species influenced by biogeographic origin. Ecol. Evol. 9, 5348–5361 (2019).

28. N. G. Smith, J. S. Dukes, Plant respiration and photosynthesis in global-scale models:\ incorporating acclimation to temperature and CO2. Glob. Chang. Biol. 19, 45–63 (2013).

29. N. G. Smith, G. Li, J. S. Dukes, Short-term thermal acclimation of dark respiration is greater in non-photosynthetic than in photosynthetic tissues. AoB PLANTS 11, plz064 (2019).

30. O. K. Atkin, I. Scheurwater, T. L. Pons, Respiration as a percentage of daily photosynthesis in whole plants is homeostatic at moderate, but not high, growth temperatures. N. Phytol. 174, 367–380 (2007).

31. O. K. Atkin, K. J. Bloomfield, P. B. Reich, M. G. Tjoelker, G. P. Asner, D. Bonal, G. Bönisch, M. G. Bradford, L. A. Cernusak, E. G. Cosio, D. Creek, K. Y. Crous, T. F. Domingues, J. S. Dukes, J. J. G. Egerton, J. R. Evans, G. D. Farquhar, N. M. Fyllas, P. P. G. Gauthier, E. Gloor, T. E. Gimeno, K. L. Griffin, R. Guerrieri, M. A. Heskel, C. Huntingford, F. Y. Ishida, J. Kattge, H. Lambers, M. J. Liddell, J. Lloyd, C. H. Lusk, R. E. Martin, A. P. Maksimov, T. C. Maximov, Y. Malhi, B. E. Medlyn, P. Meir, L. M. Mercado, N. Mirotchnick, D. Ng, Ü. Niinemets, O. S. O’Sullivan, O. L. Phillips, L. Poorter, P. Poot, I. C. Prentice, N. Salinas, L. M. Rowland, M. G. Ryan, S. Sitch, M. Slot, N. G. Smith, M. H. Turnbull, M. C. VanderWel, F. Valladares, E. J. Veneklaas, L. K. Weerasinghe, C. Wirth, I. J. Wright, K. R. Wythers, J. Xiang, S. Xiang, J. Zaragoza-Castells, Global variability in leaf respiration in relation to climate, plant functional types and leaf traits. New Phytol 206, 614 –636 (2015).

32. O. K. Atkin, M. G. Tjoelker, Thermal acclimation and the dynamic response of plant respiration to temperature. Trends Plant Sci 8, 343–351 (2003).

33. P. Ciais, C. Sabine, G. Bala, L. Bopp, V. Brovkin, J. Canadell, A. Chhabra, R. DeFries, J. Galloway, M. Heimann et al., Carbon and other biogeochemical cycles. In: Climate Change 2013: The Physical Science Basis. Contribution of Working Group I to the Fifth Assessment Report of the Intergovernmental Panel on Climate Change. Cambridge, UK: Cambridge University Press, 465–570 (2014).

34. P. B. Reich, K. M. Sendall, A. Stefanski, X. Wei, R. L. Rich, R. A. Montgomery, Boreal and temperate trees show strong acclimation of respiration to warming. Nature 531, 633 – 636 (2016).

35. P. Friedlingstein, M. O’Sullivan, M. W. Jones, R. M. Andrew, D. C. E. Bakker, J. Hauck, P. Landschützer, C. L. Quéré, I. T. Luijkx, G. P. Peters, W. Peters, J. Pongratz, C. Schwingshackl, S. Sitch, J. G. Canadell, P. Ciais, R. B. Jackson, S. R. Alin, P. Anthoni, L. Barbero, N. R. Bates, M. Becker, N. Bellouin, B. Decharme, L. Bopp, I. B. M. Brasika, P. Cadule, M. A. Chamberlain, N. Chandra, T.-T.-T. Chau, F. Chevallier, L. P. Chini, M. Cronin, X. Dou, K. Enyo, W. Evans, S. Falk, R. A. Feely, L. Feng, D. J. Ford, T. Gasser, J. Ghattas, T. Gkritzalis, G. Grassi, L. Gregor, N. Gruber, Ö. Gürses, I. Harris, M. Hefner, J. Heinke, R. A. Houghton, G. C. Hurtt, Y. Iida, T. Ilyina, A. R. Jacobson, A. Jain, T. Jarníková Jersild, F. Jiang, Z. Jin, F. Joos, E. Kato, R. F. Keeling, D. Kennedy, K. K. Goldewijk, J. Knauer, J. I. Korsbakken, A. Körtzinger, X. Lan, N. Lefèvre, H. Li, J. Liu, Z. Liu, L. Ma, G. Marland, N. Mayot, P. C. McGuire, G. A. McKinley, G. Meyer, E. J. Morgan, D. R. Munro, S.-I. Nakaoka, Y. Niwa, K. M. O’Brien, A. Olsen, A. M. Omar, T. Ono, M. E. Paulsen, D. Pierrot, K. Pocock, B. Poulter, C. M. Powis, G. Rehder, L. Resplandy, E. Robertson, C. Rödenbeck, T. M. Rosan, J. Schwinger, R. Séférian, T. L. Smallman, S. M. Smith, R. Sospedra-Alfonso, Q. Sun, A. J. Sutton, C. Sweeney, S. Takao, P. P. Tans, H. Tian, B. Tilbrook, H. Tsujino, F. Tubiello, G. R. van der Werf, E. van Ooijen, R. Wanninkhof, M. Watanabe, C. Wimart-Rousseau, D. Yang, X. Yang, W. Yuan, X. Yue, S. Zaehle, J. Zeng, B. Zheng, Global Carbon Budget 2023. doi: 10.5194/essd-2023-409 (2023).

36. P. M. Cox, R. A. Betts, C. D. Jones, S. A. Spall, I. J. Totterdell, Acceleration of global warming due to carbon-cycle feedbacks in a coupled climate model. Nature 408, 184– 187 (2000).

37. S. P. Harrison, W. Cramer, O. Franklin, I. C. Prentice, H. Wang, Å. Brännström, H. Boer, U. Dieckmann, J. Joshi, T. F. Keenan, A. Lavergne, S. Manzoni, G. Mengoli, C. Morfopoulos, J. Peñuelas, S. Pietsch, K. T. Rebel, Y. Ryu, N. G. Smith, B. D. Stocker, I. J. Wright, Eco-evolutionary optimality as a means to improve vegetation and land-surface models. N. Phytol. 231, 2125–2141 (2021).

38. S. Martorell, A. Diaz-Espejo, H. Medrano, M. C. Ball, B. Choat, Rapid hydraulic recovery in Eucalyptus. Plant, Cell Environ. 37, 617–626 (2014).

39. S. Tan, H. Wang, I. C. Prentice, K. Yang, Land-surface evapotranspiration derived from a first-principles primary production model. Environ. Res. Lett. 16, 104047 (2021).

40. S. Tan, H. Wang, I. C. Prentice, K. Yang, R. L. B. Nóbrega, X. Liu, Y. Wang, Y. Yang, Towards a universal evapotranspiration model based on optimality principles. Agric. For. Meteorol. 336, 109478 (2023).

41. S. Manzoni, P. Čapek, P. Porada, M. Thurner, M. Winterdahl, C. Beer, V. Brüchert, J. Frouz, A. M. Herrmann, B. D. Lindahl, S. W. Lyon, H. Šantrůčková, G. Vico, D. Way, Reviews and syntheses: Carbon use efficiency from organisms to ecosystems – definitions, theories, and empirical evidence. Biogeosciences 15, 5929–5949 (2018).

42. S. Vicca, S. Luyssaert, J. Peñuelas, M. Campioli, F. S. Chapin, P. Ciais, A. Heinemeyer, P. Högberg, W. L. Kutsch, B. E. Law, Y. Malhi, D. Papale, S. L. Piao, M. Reichstein, E. D. Schulze, I. A. Janssens, Fertile forests produce biomass more efficiently. Ecol. Lett. 15, 520 –526 (2012).

43. T. Thum, S. Caldararu, J. Engel, M. Kern, M. Pallandt, R. Schnur, L. Yu, S. Zaehle, A new model of the coupled carbon, nitrogen, and phosphorus cycles in the terrestrial biosphere (QUINCY v1.0; revision 1996). Geosci. Model Dev. 12, 4781–4802 (2019).

44. X. Zhou, M. Yang, Z. Liu, P. Li, B. Xie, C. Peng, Dynamic allometric scaling of tree biomass and size. Nat. Plants 7, 42–49 (2021).

45. Y. Luo, Terrestrial Carbon–Cycle Feedback to Climate Warming. Annu. Rev. Ecol., Evol., Syst. 38, 683–712 (2007).

46. Y. Ren, H. Wang, S. P. Harrison, I. C. Prentice, O. K. Atkin, N. G. Smith, G. Mengoli, A. Stefanski, P. B. Reich, Reduced global plant respiration due to the acclimation of leaf dark respiration coupled with photosynthesis. N. Phytol. 241, 578–591 (2024).

