## Supplementary Information for "Thermal acclimation of stem respiration reduces global carbon burden"

#### 1. Materials and Pre-processing

##### 1.1 Global Stem Respiration Dataset (GSRD)

Our Global Stem Respiration Dataset includes: (a) existing datasets (TRY database, which includes the Functional Ecology of Trees (FET) dataset, ECOCRAFT dataset, Global Respiration Dataset, and Tropical Respiration Dataset), (b) data digitized from publications (journal articles and book chapters from 1966 to present), (c) data provided by colleagues listed in Acknowledgments, and (d) data provided by coauthors of this article (Table S1). Our data compilation is built from individual replicates; i.e., built from data providing the species, stem respiration rate, other correlated traits and site information, to which we could reasonably assign geographic coordinates, and thus climate data, for each sample. To ensure all the data were processed in the same way, the records were screened according to the criteria described in Table S3. The data covers 96 sites worldwide, 212 species, and a total of 5275 observations, after screening. The climate space represented in our dataset includes the cold temperate zone, the temperate zone, and the tropical zone. Average temperature during the growing season (mGDD5: the average value when the daily temperature is greater than 5 ℃) ranges from 9.1 to 27.2 ℃. The range of alpha_p (AET/PET: characterizing aridity) is 0.21 to 0.82 (Figure S1).

Stem respiration was defined as the rate of $CO_2$ emission from the stem surface of a woody plant. Some measurements were made in the field and some measurements were made in the laboratory. Most studies measured stem respiration in the field (here referred to as field data)with a chamber attached to the surface of the trunk and connected to an infrared gas analyzer (IRGA) like LICOR-6400. Field data from 92 sites were obtained either from the TRY Dataset or digitized data from references. Stem diameters ranged from 10.0 to 143.0 cm, with a mean value of 25.9 ± 12.0 cm, when reported. We standardized measurements from area-based to mass-based respiration rates using living cell fractions and wood density when necessary, to reduce the effects of dimensions, according to equation 1. Based on this ,we did correction of temperature to standardize to 25 °C. The respiration rate at a standard temperature (such as 0°C or 25°C) is generally used to compare across different environments, and can be obtained using a fixed $-Q_{10}$ equation (Atkin et al., 2002) following equation 2.

$$r_{s_{mass}} = \frac{4 r_{s_{area}}}{\rho_s \times D \times SR} \#(1)$$

where $r_{s\_mass}$ and $r_{s\_area}$ is mass–based and surface–area–based sapwood respiration rate respectively (mol $CO_2$ $g^{-1}$ $s^{-1}$ and mol $CO_2$ $m^{-2}$ $s^{-1}$). $\rho_s$ is the wood density (g $m^{-3}$); D is the diameter of the stem at the measuring point, i.e. breast height ~1.3m ; SR is sapwood ratio, which is the ratio of sapwood to stem at the cross section (%).

$$r_{Tref} = r \times Q_{10}^{\frac{Tref-T}{10}} \#(2)$$

where $r_{Tref}$ is the respiration rate at the reference temperature (mol $CO_2$ $g^{-1}$ $s^{-1}$), r is the respiration rate at the measured temperature (mol $CO_2$ $g^{-1}$ $s^{-1}$), $Q_{10}$ is the thermal sensitivity of respiration, and T and $T_{ref}$ are the measuring temperature and reference temperature, respectively. We used a fixed $Q_{10}$ of 2.2 (Acosta et al.,2007). However, tests in which the $Q_{10}$ was varied from

1.6 to 2.8 (Cavaleri et al., 2006; Ryan et al., 1994; Darenova et al., 2019, 2020; Acosta et al., 2007) showed that this variation had no significant impact on the results (Table S5).

Our data set also includes 81 laboratory measurements on species from the other 4 sites (ca. 1.53% of data set). Our coauthors used core or branches removed from the field and transported back to the laboratory for mass-based sapwood respiration measurements, also using an infrared gas analysis (IRGA) machine. Although the total sample size of the laboratory measurement data is small, it has the advantages of including unified measurement units and measurement sites, and small data errors. In contrast, the field data is relatively complex, with inconsistent measurement locations and units. However, it has the advantages of a larger data volume, higher species coverage, and a wider distribution of sample sites. There was a significant difference between the laboratory and field stem respiration rate (Table S2), with laboratory measurements being approximately 6 times higher than field measurements. The mean stem respiration rate ($r_{s25}$) at standard temperature for the lab data was 2.97 nmol $CO_2$ $g^{-1}$ $s^{-1}$, with a standard error of 1.16, while the mean respiration rate for field measurements was 0.49 nmol $CO_2$ $g^{-1}$ $s^{-1}$, with a standard error of 0.0078. This difference is probably caused by some $CO_2$ leakage from the sap flow rather than emission from the bark (Levy et al. 1999; Wang et al., 2004) because bark partially limits CO2 diffusion. Although this is about stem respiration, but the bark has a limit to emission, so if condition permits, we measured sapwood respiration, which is the most important living cells here. Wang et al. (2004) compared the two measurement methods and recommended treating these two types of data separately; we have adopted this suggestion.

##### 1.1.2. Temporal stem respiration data

We also analyzed information from time series records on stem respiration. Lavigne and Ryan (1994) measured stem respiration during the 1994 growing season at eight sites with contrasting climates. We obtained data from one site near the northern boundary of the boreal region, close to Thompson, Manitoba (55.90°N, 98.75°W) from the TRY dataset, where measurements were made between May to September 1994. The species at this site included black spruce, aspen and jack pine (Gower et al., 1997).

##### 1.1.3. Warming experiment data

The warming experiment was conducted by Smith et al. (2019). They examined the thermal acclimation of stem respiration in individuals of five different woody species acclimated to five temperatures: 15, 20, 25, 30, 35 ℃. They also measured the instantaneous tissue temperature response curves (at 14, 23, 32, 41, ~50 ℃) on each individual following a 7-day acclimation period. We selected woody saplings only, with a single specimen acclimated to each temperature.

##### 1.1.4. Auxiliary trait-related data

We categorized species into plant functional types (PFTs), according to permanence of above-ground live biomass (we removed all herbaceous), leaf longevity (evergreen vs. deciduous) and leaf structure (broad-leaved vs. needle-leaved). This yields four PFTs: evergreen broad-leaved, deciduous broad-leaved, deciduous needle-leaved and evergreen needle-leaved. We assigned each species to a PFT using information from the World Flora Online (https://www.worldfloraonline.org/), the Flora of China (http://www.iplant.cn/foc) and the Plant List (http://www.theplantlist.org/).

We included information on wood density for each species, which represents the dry mass of wood per unit volume. Wood density is an important trait for estimating stored biomass and carbon content per unit volume of tree stem (Chave et al. 2009). Wood density data were obtained from the TRY data set and the African Wood Density Database (https://apps.worldagroforestry.org/treesnmarkets/wood/index.php). For each species, we found the variance of wood density within species is from 0 to 0.02 g $cm^{-3}$, suggesting this trait is conservative within species. We used the mean value of each species in our analyses.

Due to the limit of data on living cell fractions in the existing data sets, we were fairly confined to figure out the stem respiration. However, Morris et al. (2019) have assembled a data set of wood parenchyma fraction for 1439 separate species, and there is a strong relationship between parenchyma fraction and site temperature (Figure S5). For parenchyma fraction is an important proxy of living cells, which is the most important in energy released from respiration (parenchyma activity being a key determinant of respiration rates). We applied this relationship to standardize the sapwood respiration to a mass-based value for all the samples in pre-processing.

Biomass data used to simulate global sapwood respiration was from the Global Biomass Data Product (GBDP: Santoro et al., 2018). This forest map has a pixel size of 1 ha and is based on satellite remote sensing observations from the year 2010 (+/− 1year). We used the estimates of above ground biomass (AGB, unit: Mg/ha), which is defined as the dry weight of stem, bark, branches and twigs from living plants (including a few shrubs here) (i.e...,. excluding dead stumps or roots). The datasets included estimates of the per-pixel uncertainty expressed as standard error in Mg/ha (available at https://doi.org/10.1594/PANGAEA.894711).

##### 1.1.5. Environmental data

Site locations (latitude, longitude) were taken from source publications or estimated from information given therein (WGS84 datum adopted as standard). Where published coordinates did not fall in the correct country or fell in water not on land (based on the climate raster layers), new coordinates were estimated (e.g., from Google Earth) based on site descriptions or simply moved to the nearest terrestrial suitable grid−cell on the Worldclim v1.3 raster layer, matching source and model elevation as best as possible.

Temperature data for the period 1901 to 2018 were extracted from CRU TS (Climatic Research Unit gridded Time Series) version 4.02, which has a spatial resolution of 0.5° latitude by 0.5° longitude over all continents except Antarctica (Harris et al., 2020). Then we calculated the mean temperature of the growing season ($mGDD_5$) as the mean of daily temperatures on days when the temperature was > 5 °C. We calculated alpha_p, the ratio of actual evapotranspiration to potential evapotranspiration using the SPLASH model (Davis et al., 2017). The inputs included elevation (m), monthly temperature (°C), monthly sunshine hours (%) and monthly precipitation (mm). These data source from CRU CL v2.0.

Data used for calculating transpiration included: (merged with data from CRU/SPLASH??) WATCH Forcing Data methodology applied to ERA5 (WFDE5) provides bias-corrected reconstructions of near-surface meteorological variables (Cucchi et al., 2020). We extracted daily mean near-surface temperature for the site near Thompson, Manitoba (55.90° N, 98.75° W), and used a universal evapotranspiration model to calculate transpiration (Tan et al., 2021,2023). Inputs include shortwave downward radiation, atmospheric pressure in the horizontal plane, atmospheric water vapor pressure deficit, and near-surface temperature at the study site with a spatial resolution of 0.5° at hourly intervals. The fraction of absorbed photosynthetically active radiation (fAPAR) was derived from GIMMS3g with an accuracy of 1/12° in half a month, and was down-scaled to daily fAPAR by linear interpolation.

##### 1.1.6. Climate Model Outputs

We simulated stem respiration using future climate projections from CMIP6 (Coupled Model Inter-comparison Project Phase 6), following two of the Shared Socioeconomic Pathways SPP126 and SSP585 during the period of 2015 – 2100. SSP126 represents a future scenario of anthropogenic policies in which there is strong and immediate action to reduce greenhouse gas emissions. SSP585 represents a business-as-usual scenario with high greenhouse gas emissions. Four Earth System Models have data for these analyses: ACCESS – ESM1 – 5, CESM2, IPSL – CM6A – LR and UKESM1 – 0 (Table S4). We extracted near-surface temperature outputs to calculate mGDD5.

#### 2. Theoretical derivation

##### 2.1 Principles of theory predicting thermal sensitivity

By hypothesizing that sapwood respires to support its major function in maintaining the hydraulic continuum as determined by the canopy transpiration demand, we assumed that:

$$R_s = a \times E_{sap} = a \times E_{canopy} \#(3)$$

where $R_s$ (μmol $CO_2$ $m^{-2}$ $s^{-1}$) is the whole plant sapwood respiration, $E_{sap}$ (μmol $H_2O$ $m^{-2}$ $s^{-1}$) is the sapflow through the sapwood tissue, and $E_{canopy}$ the canopy – scale transpiration demand. To maintain the hydraulic continuum and lower the risk of embolism, $E_{sap}$ equals $E_{canopy}$. a is a cost factor representing the respiration rate required for supporting a unit transpiration rate.

$R_s$ is a product of the respiration rate of unit sapwood mass ($r_s$, nmol $CO_2$ $g^{-1}$ $s^{-1}$) and the total sapwood biomass ($M_s$, g C). This cost factor should, in principle, be influenced by temperature: water becomes less viscous with increasing temperature and thus the incurred cost for supporting a given transpiration rate is lower. After considering the thermal response, we obtain:

$$r_{s25} \times f(T) \times M_s = a_{25} \times \eta(T) \times E \#(4)$$

where $r_{s25}$ is mass – unit sapwood respiration rate at the reference temperature of 25°C and $a_{25}$ is the cost under this temperature. f(T) describe the kinetic thermal response by the fixed – Q10 equation (formula 6). And η(T) express the control of temperature as follows:

$$\eta(T) = e^{0.0226(25-T)} \#(5)$$

$$f(T) = Q_{10}^{\frac{T-T_{ref}}{10}} \#(6)$$

By taking logarithms on both sides of the equation, we derive the linear equation as:

$$\ln r_{s25} + 0.1 \times \ln Q_{10} + \ln M_s = \ln a_{25} + 0.0226(25 - T) + E \#(7)$$

then we obtain the general formula of $r_{s25}$:

$$\ln r_{s25} = \ln a_{25} - 0.1(T - 25) + \ln E + \ln M_s \#(8)$$

This formula can be expressed in two specific forms. The first one is assuming that $r_s$ is proportional to both E and the cost factor at a time scale when $M_s$ almost remains unchanged:

$$\ln r_{s25} = \ln a_{25} - 0.1(T - 25) + \ln E + Constant\#(9)$$

When variations in $M_s$ are too large to be ignored, as happens when considering spatial pattern, metabolic theory suggests that this variation would be canceled out through variations in E at the same time scale. Consequently, we expect $r_s$ to be proportional exclusively to the cost factor, whereas its thermal acclimation behavior remains as the short time scale:.

$$\ln r_{s25} = \ln a_{25} - 0.1(T - 25) + Constant\#(10)$$

In this case, $r_s$ at the growth temperature acclimates ($r_{s.gt}$, nmol $CO_2$ $g^{-1}$ $s^{-1}$) to track the thermal response of the cost factor as determined by water viscosity. This prediction means $r_{s.gt}$ declines with temperature by 2.3% $K^{-1}$.

#### 2.2. Sapwood respiration fraction based on EEO theory

Based on previous theory (Farquhar et al., 1980; Prentice et al., 2014; Wang et al., 2017), we can apply Fick's law to both transpiration and assimilation, and then assuming that leaves minimise the summed unit costs of transpiration and carboxylation, we can express the carbon cost of transpiration and assimilation as follows:

$$\frac{E}{A} = \frac{1.6D}{c_a(1-\chi)}\#(11)$$

$$\frac{V_{cmax}}{A} = \left(\chi + \frac{k}{c_a}\right)\frac{1}{\chi}\#(12)$$

Where E is transpiration, $V_{cmax}$ is the maximum carboxylation rate, and A is photosynthesis carbon assimilation. D is vapour pressure deficit, and ca represents ambient carbon dioxide situation. k is a function of oxygen conductance and represent the Michaelis – Menten coefficients of Rubisco. $\chi$ here is a key parameter that is determined by environmental temperature, altitude, humidity and radiation. By linking respiration with carbon cost, we get the following:

$$\frac{R_s}{A} = a\frac{E}{A} = a\frac{1.6D}{c_a(1-\chi)}\#(13)$$

$$\frac{R_d}{A} = b\frac{V_{cmax}}{A} = b\left(\chi + \frac{k}{c_a}\right)\frac{1}{\chi}\#(14)$$

$$\frac{R_{ag}}{A} = \frac{R_s}{A} + \frac{R_d}{A} = b\left(1 + \frac{k}{c_a} \times \frac{1}{\chi^2}\right)\#(15)$$

where $R_s$, $R_d$ and $R_{ag}$ represent the whole sapwood respiration, dark leaf respiration and total above – ground respiration. b is about 0.015 (Wang et al., 2020), and $\chi_0 = \frac{\xi}{\xi + \sqrt{D}}$ according the Prentice et al. (2014). The ratio of sapwood respiration to leaf respiration is then given by:

$$\frac{R_s}{R_d} = \frac{\kappa(1-\chi)}{\kappa\chi + \chi^2}\#(16)$$

where $\kappa = \frac{k}{c_a}$ , and under standard environmental conditions. Here we use 25°C for temperature,

0 m for elevation, 1 atm for atmosphere pressure and 400 ppm for $CO_2$. We used the rpmodel (Stocker et al., 2020) to calculate this ratio, which was about 22.9%~31.6%.

### 3. Methods

#### 3.1. Statistical analysis

The sapwood respiration rate data were log−transformed. The strongly right−skewed site−transpiration was also log−transformed for analysis. To compare between in field and in laboratory data, we employed Analysis of Variance (ANOVA) to assess the variation between different measurement groups.

We determined the time-scale of seasonal acclimation using $R^2$ and Variance Inflation Factors (VIF), where the highest $R^2$ and low VIF indicates the best model with low collinearity (Figure S7). The calculations were performed using the R package *car*.

Errors-in-variables (EIV) regression is a standard method for consistent estimation in linear models with noisy data covariates (Lockwood and McCaffrey 2014, 2017; Rabe−Hesketh et al., 2003). We used the R package *eivtools* for analysis of the relationship between $r_{s25}$ or $r_{s.gt}$ and $mGDD_5$ both globally and seasonally. We calculated the reliability (which is 100% minus the ratio of standard deviation and the mean value) at the global scale and seasonal scale separately. At the global scale, the time mismatch was estimated by differences over 30 years, and the spatial mismatch was calculated by comparison with the CRU CLv2.0 data. After applying error transfer equations, the reliability of global $mGDD_5$ was estimated as 0.96. At the seasonal scale, we require the reliability of temperature and transpiration. The uncertainty of temperature was estimated from the time mismatch over 3 days and the spatial mismatch was estimated by comparison with the CRU CLv2.0 data. While the uncertainty of transpiration gave extra consideration to the β parameter (the ratio of the unit costs of maintaining carboxylation and transpiration capacities) in the p−model, which contribute the most uncertainty in sensitivity analysis (Wang et al., 2017). The overall reliability of temperature and transpiration is 0.92 and 0.89 respectively.

Analysis on warming experiment data used ORL with R package *lme4*. Regression analysis was carried on site-mean values, calculated across all species and samples, and for data where the measurement temperature was 25±1°C (Table S6, S7). For the warming experiment, we conducted regression analyses separately for individual species and for the entire dataset (Table S8). We also conducted regression analysis for all measurements on *Pinu*s in the global dataset (Table S9). Figure plotting was done using the R packages *visreg* and *ggplot2*.

#### 3.2. Modelling and simulation

##### 3.2.1. A simple model of sapwood respiration at ecosystem scale

To obtain an overall estimation of sapwood respiration, we upscaled individual-scale mass-based $r_s$ to the ecosystem scale. We developed a simple model to simulate global sapwood respiration taking thermal acclimation into account:

$$F_s = \sum_{i=1}^{n} R_s \times Area \times Time_{growth} = \sum_{i=1}^{n} r_{s25} \times f(T) \times AGB \times SR \times Area \times Time_{growth} \#(17)$$

where $F_s$ is the annual carbon dioxide emission by sapwood respiration globally (unit: Pg $CO_2$). $R_s$ is land-area-based sapwood respiration (unit: nmol $CO_2$ $m^{-2}$ $s^{-1}$), and $r_{s25}$ is mass-based sapwood respiration at a reference temperature of 25 °C (unit: nmol $CO_2$ $g^{-1}$ $s^{-1}$), where we introduced thermal acclimation. f(T) follows fixed − $Q_{10}$ equation, representing instantaneous thermal response, where T is $mGDD_5$. AGB (above − ground biomass, unit: g $ha^{-1}$), SR (sapwood ratio, unit:%) and Area (pixel actual area, unit: $m^2$) are used to calculate the sapwood mass of each pixel. Time_growth is defined as the period when the mean temperature is above 5 °C (unit: °C).

We obtained the value of $r_{s25}$ by calibration using data from the GSRD where the measurement temperature was 25±1°C. And by fixing the sensitivity of 0.1, we did regression between $\ln r_{s25}$ and $mGDD_5$. That means for natural situation, the referenced sapwood respiration can be captured by $mGDD_5$ and predict the $r_{s25}$ worldwide. The generalized expression of $r_{s25}$ is as follows,

$$ln\, r_{s25} =- 0.1(mGDD_5 - 25) + C\#(18)$$

From this calibration, the estimated value of C is −1.843, and the standard error of the $r_{s25}$ equals that of intercept of 0.039 (Figure S6).

##### 3.2.2. Simulation of sapwood respiration in 2010, as an example

We applied this simple model using above-ground biomass data around 2010 from the GBDP and daily mean temperature from CRU. All data was standardized to a spatial resolution of 0.5° to run the model. The global $CO_2$ emission was calculated through summation. We used quadratic mean value to calculate SE of sapwood respiration. The model uncertainty came from $r_{s25}$, biomass and sapwood ratio approximation. The standard error of $r_{s25}$ and sapwood ratio is 0.039 and 0.19 separately. The GBDP biomass data has a standard error per grid cell; this introduced the most uncertainty.

$$SE_{R_s} = \sqrt{{SE_{AGB}}^2 + {SE_{SR}}^2 + {SE_{r_{s25}}}^2}\#(19)$$

Our estimate of the annual $CO_2$ released by sapwood respiration globally is 11.20 ± 5.88 Pg

C.

##### 3.2.3. Simulation of future sapwood respiration under different scenarios

To estimate the effect of considering the thermal acclimation of sapwood respiration, we calculated the carbon emission reduction over the period 2015-2100 with and without introducing acclimation ($F_{s.re}$), using outputs from four models and two scenarios The reduction caused by considering thermal acclimation (Fs.re) can be expressed as follows:

$$F_{s.re} = \sum_{i=1}^{n} \left[ (r_{s25} - r_n) \times f(T) \times AGB \times SR \times Area \times Time_{growth} \right] \#(20)$$

where $F_{s.re}$ represents the carbon emission reduction compared to non-acclimated respiration in each year. $r_{s25}$ is the referenced mass-based sapwood respiration calculated by the model. $r_n$ is the referenced sapwood respiration from the modern parameter calibration and is kept constant in the future simulations. The temperature used to calculate $r_{s25\_a}$, f(T) and $Time_{growth}$ was extracted from CMIP6 models.

We compared our model with results from CLM5, which takes thermal acclimation into account as:

$$r_{s25_{clm5}} = r \times 10^{(-0.00794 \times (mGDD_5 - 25))} \#(21)$$

where $r_{s25\_clm5}$ means referenced sapwood respiration considering thermal acclimatio, r is a PFT-specific parameter. The natural-log sapwood respiration in CLM5 decreases with temperature by 1.83 % $K^{-1}$, about 2/11 of our theoretical prediction. Applying this equation to the future SSP126 and SSP585 scenarios, would result in a release of ca. 3.6 and 7.8 Pg C additional $CO_2$ by 2100 compared to our simulations.

### Supplementary Figures and Tables

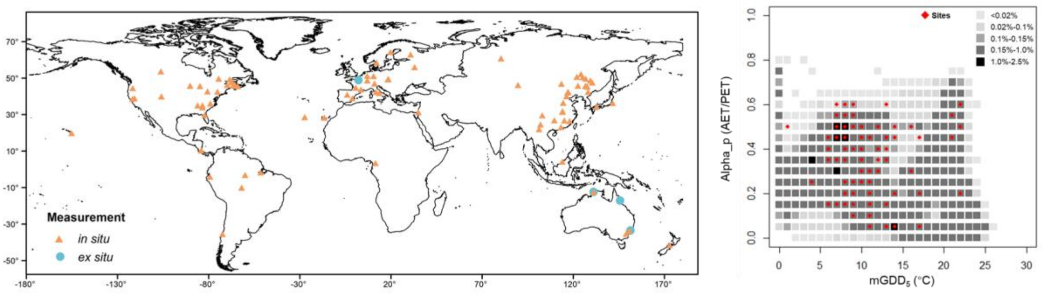


**Fig. S1. Sites.** The map shows the location of sites used for field (*in situ*) and laboratory (*ex situ*) measurements of sapwood respiration. The right-hand plots shows the location of the samples (red points) in climate space, defined by alpha_p (AET/PET) and mean growing season temperature (mGDD5). The grey shading indicates the frequency of each climate globally.

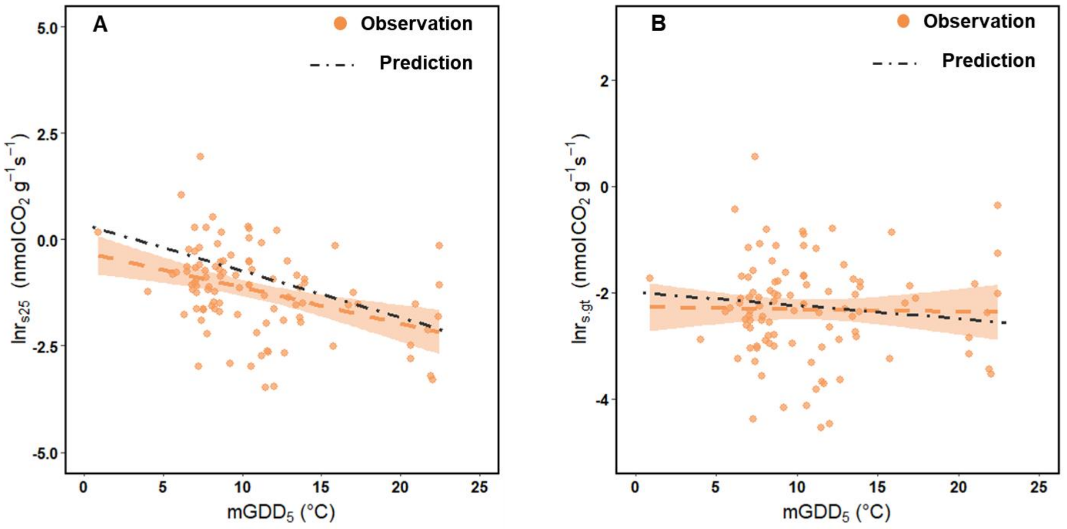


**Fig.S2. Global trends of sapwood respiration and temperature based on site-mean values.**

$r_{s25}$ represents sapwood respiration standardized to the reference temperature of 25 ℃, $r_{s.gt}$ represents sapwood respiration standardized to the mean growing temperature, and $mGDD_5$ is the mean temperature of the growing season defined as days when the temperature is >5℃.

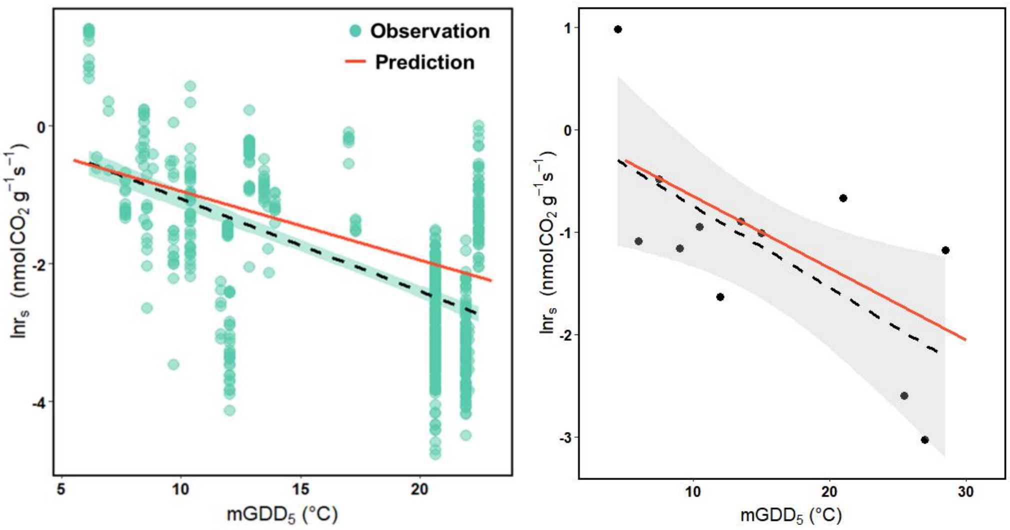


**Fig.S3. Global trends of sapwood respiration based on samples with no temperature standardization.** The measurements of rs were conducted within 1℃ of 25℃. The left panel displays the raw data, while the right panel shows the data after temperature has been divided into 20 bins.

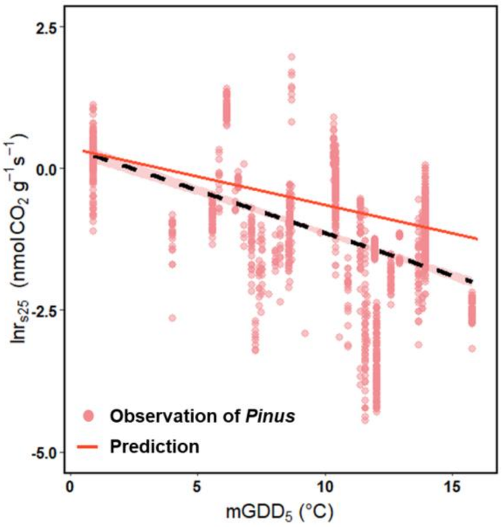


**Fig.S4. Global trends in sapwood respiration and temperature based on measurements of *Pinus*.** $r_{s25}$ represents sapwood respiration standardized to the reference temperature of 25 ℃, and $mGDD_5$ is the mean temperature of the growing season defined as days when the temperature is >5℃.

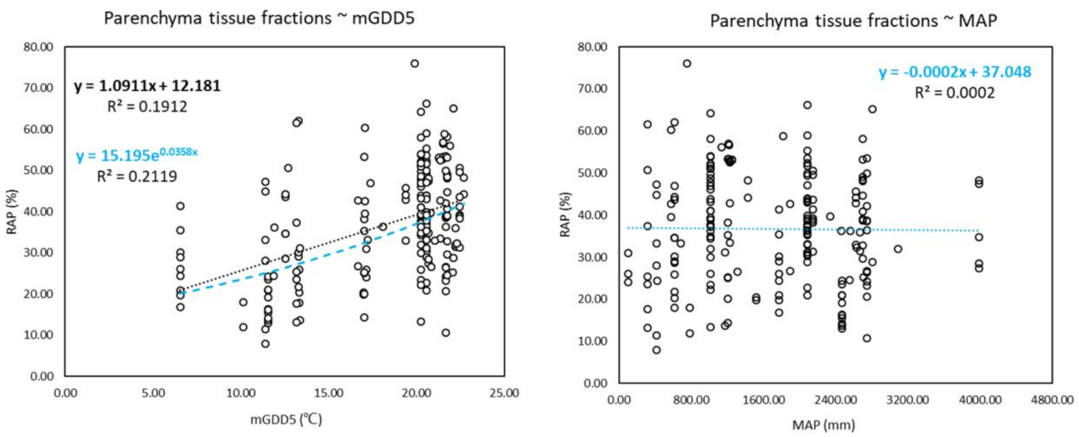


**Fig.S5.** The relationship between sapwood ratio environmental factors, specifically the mean temperature of the growing season defined as days when the temperature is >5°C ($mGDD_5$) and mean annual precipitation (MAP)

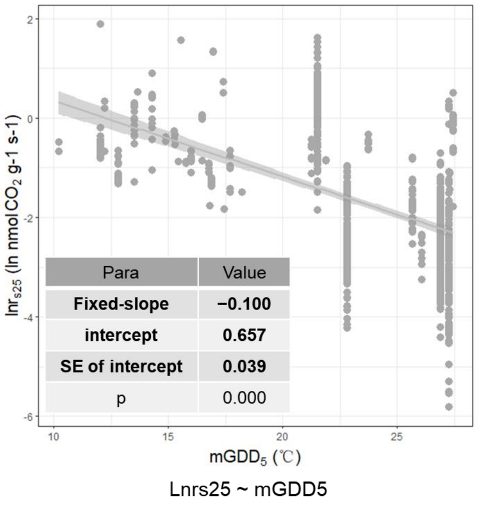


**Fig.S6. Parameter calibration of sapwood respiration model.** The calibration was based on samples measured at temperatures of 25±1°C. The linear regression used a fixed slope of −0.1 (theoretical predicted value) to estimate the intercept and its standard error, representing the parameter's uncertainty.

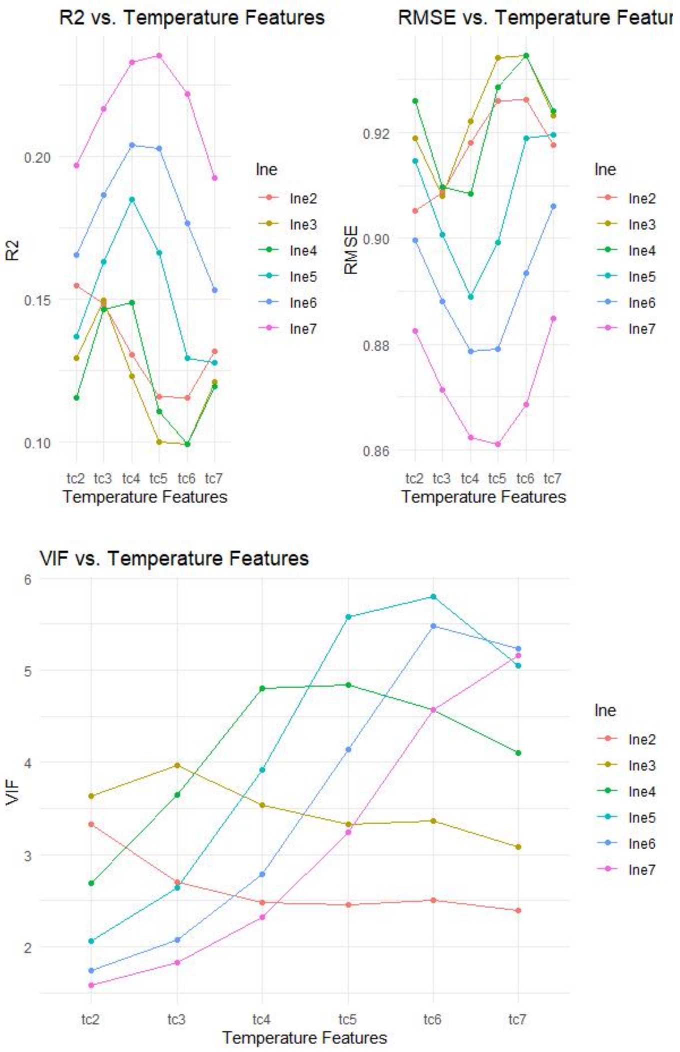


**Fig.S7. Identification of acclimation time-scale.** The plots show the $R^2$, root mean square error (RMSE) and variance inflation factor (VIF) of different time-scale combinations of temperature and transpiration, where tc2, tc3, tc4, tc5, tc6 and tc7 are the mean temperature over the 2, 3, 4, 5, 6 and 7 days before sapwood respiration was measured and the transpiration was natural-log transformed.

**Table S1 Sites location and origins of sapwood respiration**

| Latitude (°) | Longitude (°) | Ref ID | Origin | Measurement Methods | Number of Samples |
|---|---|---|---|---|---|
| 42.4 | 128.1 | 1 | digitised | In field | 201 |
| 26.75 | 115.07 | 2 | digitised | In field | 71 |
| 26.47 | 117.95 | 3 | digitised | In field | 24 |
| 26.47 | 117.95 | 4 | digitised | In field | 96 |
| 26.85 | 109.6 | 5 | digitised | In field | 132 |
| 26.75 | 115.07 | 6 | digitised | In field | 187 |
| 45.33 | 127.57 | 7 | digitised | In field | 61 |
| 21.93 | 101.27 | 8 | digitised | In field | 72 |
| 45.4 | 127.67 | 11 | digitised | In field | 270 |
| 45.4 | 127.67 | 12 | digitised | In field | 36 |
| 36.42 | 114.5 | 13 | digitised | In field | 30 |
| 42.315 | 117.25 | 14 | digitised | In field | 10 |
| 45.4 | 127.67 | 16 | digitised | In field | 39 |
| 24.53 | 102.03 | 17 | digitised | In field | 66 |
| 45.33 | 127.57 | 18 | digitised | In field | 12 |
| 42.03 | 116.85 | 19 | digitised | In field | 36 |
| 23.13 | 113.28 | 21 | digitised | In field | 72 |
| 29.57 | 102.99 | 22 | digitised | In field | 4 |
| 45.72 | 126.63 | 25 | digitised | In field | 6 |
| 40.03 | 116.33 | 27 | digitised | In field | 31 |
| 26.75 | 115.07 | 28 | digitised | In field | 34 |
| 48.67 | 7.08 | 33 | original data collector | In field | 16 |
| 49.5 | 18.53 | 35 | original data collector | In field | 13 |
| 56 | 33 | 36 | digitised | In field | 167 |
| −3.12 | −60.02 | 44 | digitised | In field | 24 |
| 38.54 | −121.8 | 46 | digitised | In field | 5 |
| 39.9 | −105.87 | 51 | original data collector | In field | 22 |
| 10.43 | −84 | 52 | original data collector | In field | 2 |
| 38.9 | −120.63 | 60 | digitised | In field | 72 |
| 47.55 | 130.42 | 67 | digitised | In field | 5 |
| 48.67 | 129.42 | 67 | digitised | In field | 5 |
| 49.62 | 126.8 | 67 | digitised | In field | 5 |
| 50.45 | 125.2 | 67 | digitised | In field | 5 |
| 50.62 | 121.95 | 67 | digitised | In field | 5 |
| 52.19 | 124.22 | 67 | digitised | In field | 5 |
| 29.73 | −82.15 | 70 | original data collector | In field | 72 |
| 45.75 | 122.58 | 70 | original data collector | In field | 1 |
| 46.17 | 89.67 | 70 | original data collector | In field | 1 |
| 46.85 | 113.48 | 70 | original data collector | In field | 1 |
| 49.69 | −74.43 | 72 | digitised | In field | 11 |
| 43.73 | 3.58 | 73 | digitised | In field | 29 |

| | | | | | |
|---|---|---|---|---|---|
| 49.5 | 18.53 | 74 | digitised | In field | 12 |
| −33.62 | 150.74 | 76 | digitised | In field | 5 |
| −4.06 | −79.25 | 79 | digitised | In field | 353 |
| 28.3 | −16.57 | 80 | digitised | In field | 41 |
| 36.32 | 141.58 | 81 | digitised | In field | 12 |
| 35 | −86 | 83 | digitised | In field | 52 |
| 58.38 | 12.15 | 84 | digitised | In field | 6 |
| 45.59 | −84.71 | 87 | digitised | In field | 112 |
| 45.8 | −90.12 | 88 | digitised | In field | 4 |
| 4.2 | 114.03 | 91 | digitised | In field | 7 |
| 36.417 | 114.5 | 96 | digitised | In field | 16 |
| 26.833 | 109.6 | 109 | digitised | In field | 12 |
| 10.26 | −84 | 115 | digitised | In field | 187 |
| 49.3 | 18.32 | 116 | digitised | In field | 132 |
| −41.58 | 172.75 | 117 | digitised | In field | 24 |
| 31.49 | 114.035 | 120 | digitised | In field | 58 |
| 36 | −79 | 121 | digitised | In field | 6 |
| 33.88 | −83.36 | 122 | digitised | In field | 50 |
| 53.63 | −106.2 | 124 | digitised | In field | 16 |
| 41.85 | 13.58 | 125 | digitised | In field | 27 |
| 42.37 | 11.8 | 126 | digitised | In field | 30 |
| 34.38 | 132.65 | 135 | digitised | In field | 131 |
| 42.66 | −80.58 | 135 | digitised | In field | 1 |
| 45.55 | −64.05 | 138 | original data collector | In field | 10 |
| 45.8 | −65.26 | 138 | original data collector | In field | 6 |
| 46.2 | −69.8 | 138 | original data collector | In field | 11 |
| 46.53 | −65.87 | 138 | original data collector | In field | 10 |
| 47.86 | −67.46 | 138 | original data collector | In field | 6 |
| 48 | −67.46 | 138 | original data collector | In field | 11 |
| 48.11 | −67.32 | 138 | original data collector | In field | 10 |
| 49.3 | −68.4 | 138 | original data collector | In field | 6 |
| 45.95 | −66.6 | 140 | original data collector | In field | 23 |
| 47 | 11 | 141 | digitised | In field | 142 |
| 44.5 | −121.62 | 142 | digitised | In field | 9 |
| 31.33 | 35.05 | 147 | digitised | In field | 22 |
| 38.95 | −1.15 | 151 | digitised | In field | 187 |
| 64.23 | 19.76 | 152 | digitised | In field | 51 |
| 42.5 | −75.167 | 155 | digitised | In field | 40 |
| −35.35 | 148.93 | 160 | original data collector | In field | 5 |
| 19.84 | −155.125 | 163 | original data collector | In field | 3 |
| 60.75 | 80.38 | 165 | digitised | In field | 11 |
| 40.867 | −4.017 | 167 | digitised | In field | 9 |
| 51.225 | 6.946 | 168 | digitised | In field | 35 |
| 34.38 | 132.65 | 170 | digitised | In field | 70 |

| | | | | | |
|---|---|---|---|---|---|
| −35.5 | −72.41 | 171 | digitised | In field | 17 |
| −33.62 | 150.74 | 173 | digitised | In field | 119 |
| 28.583 | −27.25 | 177 | digitised | In field | 36 |
| 45.38 | 127.53 | 182 | digitised | In field | 506 |
| 31.82 | 114.07 | 185 | digitised | In field | 36 |
| −3.12 | −60.02 | 195 | digitised | In field | 10 |
| −10.08 | −61.92 | 113TRY | original data collector | In field | 28 |
| 3.38 | 11.5 | 113TRY | original data collector | In field | 57 |
| 44.7 | 0.77 | 197TRY | original data collector | In field | 1 |
| −43 | 170.3 | 198TRY | original data collector | In field | 6 |
| 48.83 | 2.17 | 199TRY | original data collector | In lab | 8 |
| 51.08 | 10.45 | 202TRY | original data collector | In field | 172 |
| 35.03 | −83.38 | 207TRY | original data collector | In field | 19 |
| 62.87 | 30.82 | 208TRY | original data collector | In field | 17 |
| −12.48 | 131.15 | 31TRY | original data collector | In field | 56 |
| 48.67 | 7.07 | 32TRY | original data collector | In field | 37 |
| −1.72 | −51.45 | 50Lucy | co−author | In field | 97 |
| 35 | −86 | 90TRY | original data collector | In field | 38 |
| −12.46 | 131.11 | Darwin | co−author | In lab | 17 |
| −17.12 | 145.62 | Qld | co−author | In lab | 41 |
| −33.58 | 151.29 | Sydney | co−author | In lab | 15 |

**Table S2. ANOVA result of in field and in laboratory sapwood respiration**

| | **Df** | **Sum Sq** | **Mean Sq** | **F value** | **Pr (>F)** |
|---|---|---|---|---|---|
| Measurement Methods | 1 | 135 | 134.91 | 123.5 | <2e−16*** |
| Residuals | 5275 | 5763 | 1.09 | | |

Signif. codes: 0 ‘***’

**Table S3 Criteria for data filtering**

| Criterion | Details |
|---|---|
| Age | Measurements on adult trees, using a minimum diameter of 10 centimeters to exclude saplings. |
| Species/Size | Only used records that provide tree diameter and name of the species. |
| Time of the year | Measurements taken within the growing season. In sites with a dry and wet season, the measurements were made in the wet season. The growing season and wet season was defined by each specific researcher |
| Measuring Height | Measurements were made at or near breast height (about 1.5 meters above the ground) to exclude vertical variation. |
| Orientations | Measurements at a given site were made on the same side of the tree trunk to avoid the impact of different trunk orientations on respiration rate. |
| Control factors | Measurements were made on woody plants in their natural growth state, without human control factors such as soil nutrient gradients, carbon dioxide fertilization, bark girdling, sparse forests and so on. |
| Temperature | Source studies provided the temperature when the respiration was measured to allow standardization to a reference temperature. |

**Table S4 Output model ensemble from CMIP6**

| ESM | LSM | Details of LSM | Reference |
|---|---|---|---|
| ACCESS−ESM1−5 | CABLE2.2.3 | Land surface models used to simulate interactions among the atmosphere, ecosystems, and land | Wang et al. (2020). |
| CESM2 | CLM5 | Emphasizes the coupling of land-atmosphere-hydrological processes, including land carbon cycling, water cycling, and ecosystem dynamics. It also includes a rich set of vegetation and soil parameterization options to adapt to various geographical conditions | Lawrence et al. (2019). |
| IPSL−CM6A−LR | ORCHIDEEV2.0 | Features high-resolution meteorological driving and detailed ecosystem processes, suitable for ecosystem and climate research | Krinner et al. (2005) |
| UKESM1−0 | JULES−ES−1.0 | It emphasizes the exchange of matter and energy processes between land surface and the atmosphere, including land carbon cycling, water cycling, and ecosystem dynamics. | Clark et al. (2011) |

**Table S5. Validating stability with varying $Q_{10}$ parameter values**

| $Q_{10}$ | Theory Prediction | Fitted slope |
|---|---|---|
| 1.6 | −0.070 | −0.099 (se=0.003) |
| 1.8 | −0.0814 | −0.106 (se=0.003) |
| 2.0 | −0.0919 | −0.112 (se=0.003) |
| 2.2 | −0.100 | −0.118 (se=0.003) |
| 2.4 | −0.110 | −0.123 (se=0.003) |
| 2.6 | −0.118 | −0.128 (se=0.003) |

**Table S6. Mathematical details of site−mean sapwood respiration (Fig.S2).**

| Measurement | Prediction | fitted coefficient | confidence intervals 2.50% | 97.50% | Intercept (mean ± SE) | R2 | p | df |
|---|---|---|---|---|---|---|---|---|
| in field | –0.1 | –0.134 | −0.142 | −0.127 | 0.289 ± 0.135 | 0.34 | <0.001 | 615 |

**Table S7. Mathematical details of sapwood respiration without temperature standardization (Fig.S3).**

| Quantity | Theoretical prediction | fitted coefficient | confidence intervals 2.50% | 97.50% | Intercept (mean ± SE) | R2 | p | df |
|---|---|---|---|---|---|---|---|---|
| lnrs25 | −0.100 | −0.083 | −0.104 | −0.062 | −0.301 ± 0.244 | .140 | <0.001 | 97 |
| lnrs.gt | −0.023 | −0.004 | −0.0253 | 0.0165 | −2.272 ± 0.244 | .000 | >0.5 | 97 |

**Table S8. Mathematical details of each species in warming experiment (Fig.3).**

| Quantity | Theoretical prediction | fitted coefficient | confidence intervals | | Intercept (mean ± SE) | R2 | p | df |
|---|---|---|---|---|---|---|---|---|
| | | | 2.50% | 97.50% | | | | |
| B.alleghaniensis | | −0.084 | −0.096 | −0.072 | 7.34 ± 0.32 | .44 | <0.001 | 57 |
| P. nigra | | −0.093 | −0.103 | −0.082 | 6.85 ± 0.28 | .49 | <0.001 | 73 |
| P. pinaster | | −0.096 | −0.109 | −0.083 | 7.04 ± 0.34 | .43 | <0.001 | 73 |
| P. pinea | −0.100 | −0.122 | −0.155 | −0.090 | 7.27 ± 0.75 | .21 | <0.001 | 50 |
| P. sylvestris | | −0.076 | −0.094 | −0.058 | 6.26 ± 0.46 | .19 | <0.001 | 72 |
| All species | | −0.088 | −0.096 | −0.081 | 6.82 ± 0.19 | .30 | <0.001 | 325 |

**Table S9. Mathematical details of sapwood respiration of global *Pinus* (Fig.S4).**

| Prediction | fitted coefficient | confidence intervals | | $R^2$ | p | df |
|---|---|---|---|---|---|---|
| | | 2.50% | 97.50% | | | |
| –0.10 | –0.15 | –0.14 | –0.13 | .253 | <0.001 | 1586 |
